# mTORC2 combats cellular stress and potentiates immunity during viral infection

**DOI:** 10.1101/2020.10.09.333500

**Authors:** Rahul K Suryawanshi, Chandrashekhar D. Patil, Alex Agelidis, Raghuram Koganti, Joshua M. Ames, Lulia Koujah, Tejabhiram Yadavalli, Krishnaraju Madavarju, Lisa M. Shantz, Deepak Shukla

**Affiliations:** Department of Ophthalmology and Visual Sciences, University of Illinois at Chicago, Chicago IL, USA; Department of Microbiology and Immunology, University of Illinois at Chicago, Chicago, IL 60612, USA; Department of Cellular and Molecular Physiology, Pennsylvania State University College of Medicine, Hershey, PA, USA

**Keywords:** mTORC2, HSV-1, Systemic infection, immune response, apoptosis

## Abstract

Herpes simplex virus type 1 (HSV-1) causes ocular and orofacial infections, which are generally well controlled by the host and nonlethal. In rare cases, HSV-1 causes encephalitis, which leads to permanent brain injuries, memory loss or even death. Host factors protect the organism from viral infections by activating the immune response. However, the factors that confer neuroprotection during viral encephalitis are unknown. Here we show that mammalian target of rapamycin complex 2 (mTORC2) is essential for the host survival of ocular HSV-1 infections *in vivo*. We found that the loss of mTORC2 causes systemic HSV-1 infection not only because of weak innate and adaptive immune responses but also due to increased ocular and neuronal cell death, which becomes lethal over time. Furthermore, we found that mTORC2 mediates cell survival channels through the inactivation of the proapoptotic factor FoxO3a. Our results demonstrate how mTORC2 potentiates host defenses against viral infections as well as implicating mTORC2 as a necessary host factor for survival. We anticipate our findings may help develop new therapeutic window for severe HSV-1 infections, such as herpes simplex encephalitis.

## Main

Herpes simplex virus-1 (HSV-1) is one of the most notorious virus infections known to mankind. An estimated 3.7 billion people under age 50 (67%) have HSV-1 infections globally. HSV-1 causes distressing ocular infections and, in severe cases, life threatening diseases like viral encephalitis which results in permanent brain damage. After primary infection, HSV-1 remains latent in neuronal tissues and reactivates under stress situations. In general, the ~74 viral proteins usurp the host assembly to make the host microenvironment permissive to virus replication. However, several host proteins combat the viral infection by inducing innate and adaptive immune responses. Host cells also activate stress responses, including programmed cell death, which limits spread of virus^1^. Although apoptosis plays an important role in antiviral defense mechanisms^2^, excessive cell death in neurons during neuronal HSV-1 infection may be detrimental to host survival^3^. This phenomenon might be one of the reasons behind severities during viral encephalitis. Exploring the role of pro-survival host factors may shed light on ways to combat severe HSV-1 infections.

mTORC2 is a protein complex known to regulate cell survival and immune signaling during viral infection^4, 5^. However, the role of mTORC2 in systemic infections has not yet been explored. To determine whether the expression of Rictor, an essential component of mTORC2, is modified during HSV-1 infection, we infected human corneal epithelial (HCE) cells at varying MOIs. By 24 hours post infection (hpi), Rictor transcripts increased in a dose-dependent manner with HSV-1 infection (Extended Data Fig. 1a). However, Rictor proteins levels briefly decline upon the onset of early viral gene expression such as ICP-0 and ICP-4 (Extended Data Fig. 1b). We hypothesized that there may exist an interplay between HSV-1 and Rictor, but there are no pharmacological inhibitors which selectively target mTORC2. Furthermore, genetic deletion of Rictoris embryonically lethal^6^. To bypass these limitations, we utilized a CreERT2 model to engineer Rictor conditional knockout mice (Extended Data Fig. 1c)^7^. Upon tamoxifen treatment, Rictor can be deleted from these mice, which henceforth referred to as iRic -/- (inducible rictor knockout) mice.

After five days of tamoxifen treatment, iRic +/+ and -/- mice were infected with HSV-1 at 1×10^5^ MOI to their corneas (Fig. 1a). Strikingly, viral infection in the iRic -/- mice is nearly lethal. While none of the iRic +/+ mice ceased at 13 dpi, 80% of the Ric -/- mice died by the same time point (Fig. 1b). We hypothesized that a systemic infection beyond the cornea led to the animal death. We used another set of iRic +/+ and -/- mice to investigate the progress of the HSV-1 infection. Eye washes were taken from the mice at 4 dpi (Fig. 1a). By 4 dpi, iRic -/- contained more virus in their eye washes (Fig. 1c). iRic -/- mice also displayed more severe corneal damage and a greater infection score (Fig. 1d, e). In addition, iRic -/- lost significantly more weight than iRic +/+ and exhibited greater quantities of viral transcripts in their blood (Fig. 1f, 1g). These results constitute evidence of a more severe, systemic infection in the iRic -/- mice.

**Figure 1.**
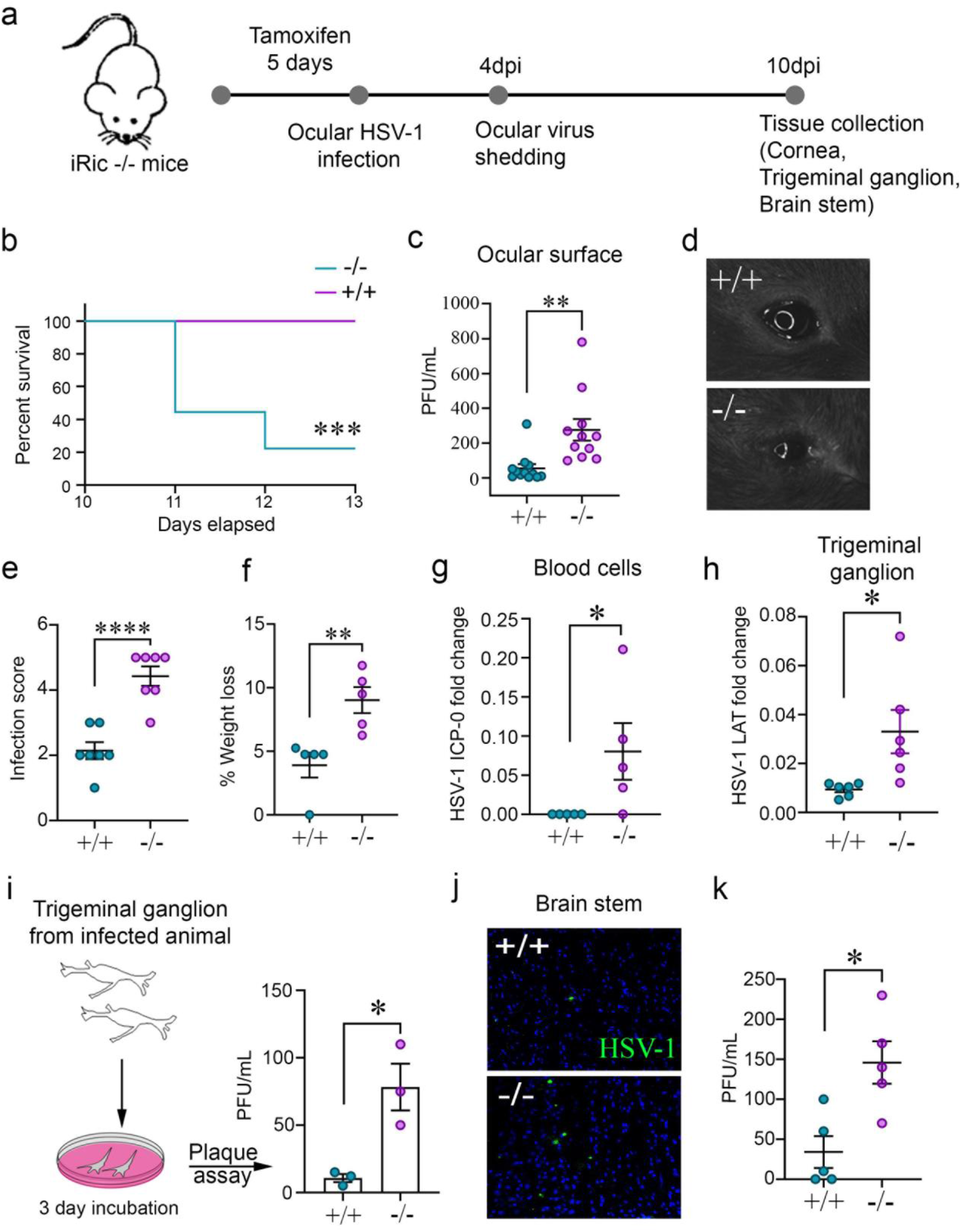
Lack of Rictor is lethal and causes systemic HSV-1 infection. **a,** Schematics of the experiment showing conditional knockout of the iRic-/- mice using Tamoxifen or mock. The mice were infected with HSV-1 analyzed for systemic infection. **b,** Survival curve showing percent survival after iRIC +/+ and -/- infection. **c,** Plaque assay showing mature virus particle formation in eyewash samples collected at 4 hpi. **d**, Representative eye image of HSV-1 infected animals at 4dpi. **e,** Graph showing image score based on visual observation at 6dpi. **f,** Graph representing percent weight loss at 12dpi. **g**,qRTPCR analysis of early viral gene isolated from blood cells at 5dpi. **g,** qRTPCR analysis of late viral gene isolated from trigeminal ganglion (TG) cells at 5dpi. **h,** To reactivate the virus in TG of infected mice, TG were cultured in vivo and the mature virus particles were analyzed with plaque assay. **i,** A representative micrograph of immunohistochemistry analysis for HSV-1 detection in brain stem at 10dpi. **j,** Graph representing mature virus particle formation in brain stem at 5dpi.

During ocular infection, HSV-1 travels from the cornea to the trigeminal ganglion (TG)^8^. qRTPCR of the TG tissue revealed an increased presence of the HSV-1 Lat transcripts, a marker for active viral replication, in the knockout mice (Fig. 1h)^9^. Furthermore, after culturing murine TGs from infected mice *ex vivo*, we observed more viral plaques in TGs isolated from iRic -/- mice (Fig. 1i). Since the TG yielded more virus in the iRic -/- mice, we wanted to investigate the viral load in the brain stem which is the next destination of HSV-1 after it reaches the TG. Both immunostaining and plaque assay results demonstrate an increased viral presence in the brain stems of the knockout mice (Fig 1j, 1k). Collectively, our data indicate that the iRic -/- mice displayed a worse HSV-1 infection in the tissues of the cornea, TG, and brain stem. Thus, HSV-1 infection in the Rictor knockout mice results in a severe, systemic infection which culminates in their death.

Given the disparities in survivability between the iRic +/+ and -/- mice during infection, we wanted to identify whether the immune response of the knockout mice was intact. Murine corneas taken at 12 dpi exhibit greater immune cell infiltration in the wild-type mice (Fig. 2a). We then checked the presence of a panel of immune cells using flow cytometry on the whole eye tissue. Here too, the iRic +/+ possessed greater numbers of helper T cells (Fig. 2b), cytotoxic T cells (Fig. 2c), active T cells (Fig. 2d), dendritic cells (Fig. 2e), and NK cells (Fig. 2f). The populations of plasmacytoid dendritic cells remained similar across the two groups (Fig. 2g). We confirmed the influx of CD8a T cells in the wild-type corneal tissue via IHC (Fig. 2h). Since the T cell populations were decreased in the iRic -/- mice, we measured whether the B cell response was attenuated as well. After infecting both groups of mice and isolating blood serum at 10dpi, we performed a serum neutralization assay (Fig. 2i). The concentration of neutralizing antibodies was far greater in the wild-type mice than the knockouts, suggesting that both humoral and cell-mediated immunity were defective in the iRic -/- mice (Fig. 2j). The impaired B cell response in absence of mTORC2 might be due to multiple reasons including downregulation of IL-7 receptor and functional follicular helper CD4+ T cell that results in IgM+ immature B cell generation^10, 11^.

**Figure 2.**
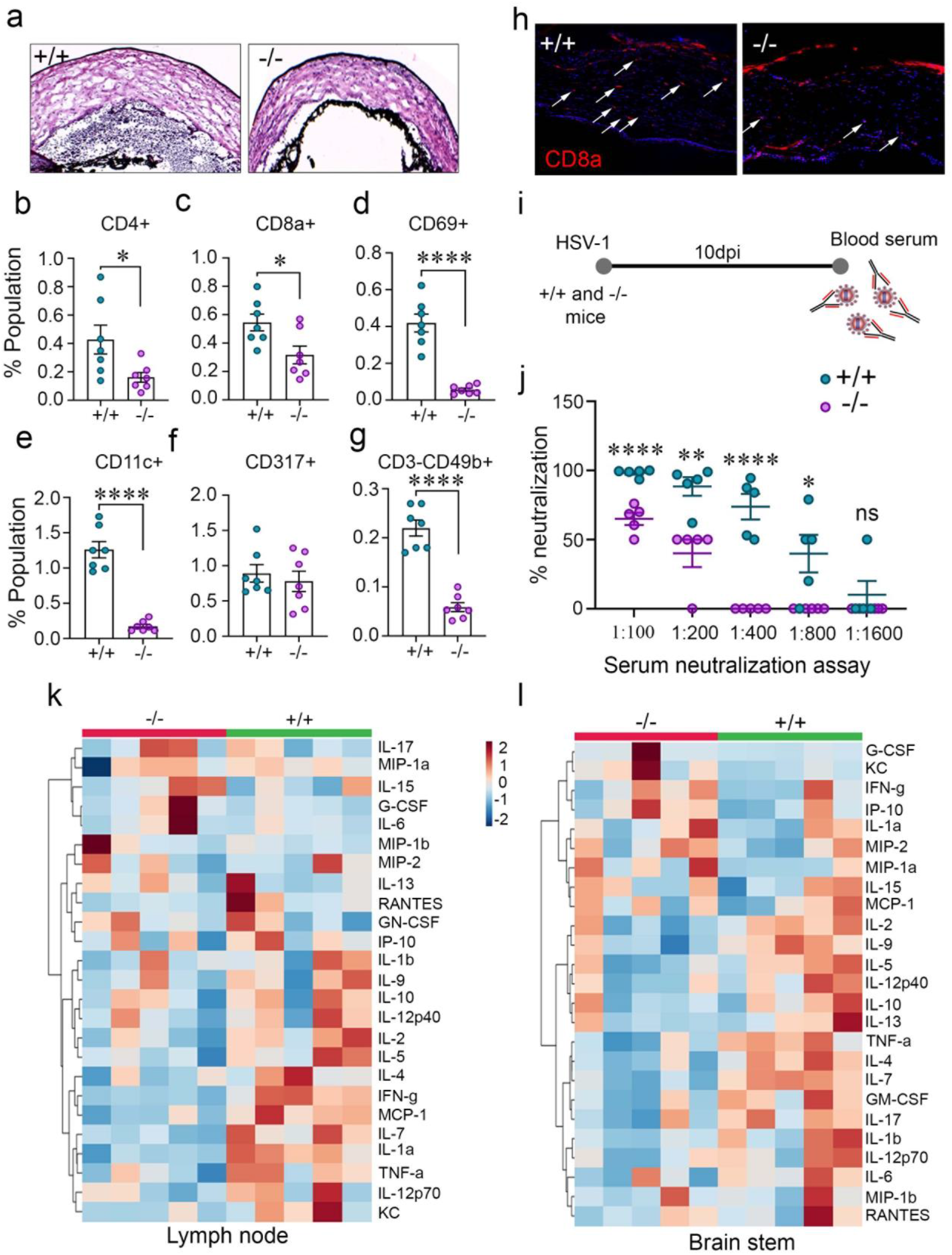
Rictor deficiency results in defective innate and adaptive immune responses. **a,** A representative micrograph for H and E staining of HSV-1 infected corneal tissue. **b-g,** Graph representing population of respective immune cells in HSV-1 infected eye at 12dpi. **H,** Immunohistochemistry staining of CD8+ cells in HSV-1 infected corneal tissue. **i,** Schematics of neutralization assay. **j,** Graph showing neutralizing antibody response from serum collected at 10dpi. **k,** Cytokine analysis of lymph node and **l,** brain stem tissue of HSV-1 infected animals at 10dpi.

As both arms of the adaptive immune response were attenuated in the Rictor knockout mice, it was possible that their activation and coordination were impaired. Using tissue isolated from cervical lymph nodes and the brain stem, we explored the cytokine profile of each group of mice using a Luminex cytokine panel. In both the tissues, the iRic -/- mice showed a weakened cytokine response compared to their wild-type counterparts (Fig. 2k, 2l). Furthermore, the types of cytokines activated by each group of mice significantly differed. In the brain stem, the pro-inflammatory cytokines IL-1α, IL-1β, and TNF-αwere up-regulated in the wild-type mice (Extended Data Fig. 2). However, the expression of the anti-inflammatory cytokines IL-2, IL-10, IL-13, IL-17 was increased in the Rictor knockout mice, both at baseline and after infection (Extended Data Fig. 2) and the results are in line with previous reports^12, 13^. These patterns suggest that mTORC2 may limit the expression of anti-inflammatory cytokines highlighting important role on mTORC2 in their homeostasis. The iRic -/- mice appear to have an intrinsically weakened immune system which cannot stimulate adaptive immune responses during infection. In agreement with our results, mTORC2 has been reported to be required for the induction of CCL5 in neurons and astrocytes during infection^5^. Collectively, these findings indicate that iRic -/- mice have deficiencies in the activation and functioning of the adaptive immune response.

mTORC2 plays important roles in balancing cell survival and apoptosis to promote homeostasis of the cell^14^. We have shown that Rictor knockout mice have defects in their immune response. During viral infection, the iRic -/- mice would not be able to rely on the adaptive immune response for protection, so we hypothesized that they may be more likely to undergo cellular apoptosis to mitigate viral spread. This shift in homeostasis towards cell death may account for the difference in survivability between the iRic +/+ and -/- mice. We performed a TUNEL stain of the corneas of infected mice at 4 dpi and observed that the knockout mice show a marked pattern of cell death along the cornea and retina, which extends to the optic nerve (Fig. 3a). In parallel, TUNEL staining of the brain stem at 10 dpi highlighted an increase in cell death in the iRic -/- mice (Fig. 3b). However, iRic +/+ showed relatively little cell death in these two tissues. The cell death in eye tissue and neurons may be attributed to impaired AKT activation by mTORC2^15, 16^.

**Figure 3.**
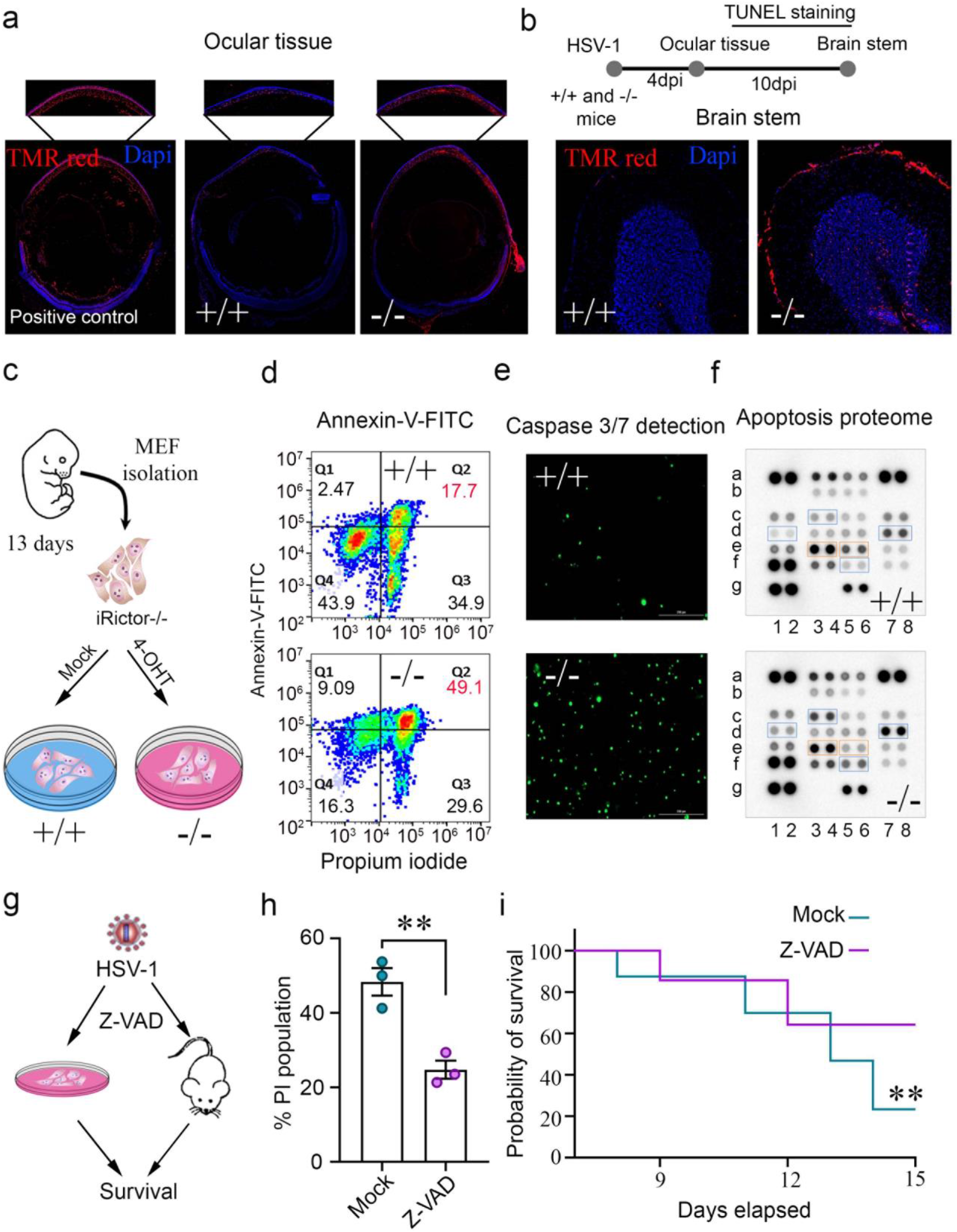
Absence of Rictor induces apoptosis. **a,** A representative micrograph of TUNEL stained whole eyeball tissue at 4dpi. **b,** Schematic and micrograph showing TUNEL staining of brain stem at 10dpi. **c,** Schematic of mouse embryonic fibroblasts (MEFs) isolation and western blot showing deletion of RICTOR by 4-hydroxitamoxifen treatment. **d,** Annexin-V-FITC and PI staining of MEFs at 12 hours post infection (hpi). Q1. Annexin V+/PI-cells indicate cells undergoing early apoptosis, Q2. Annexin V+/PI+ indicate late apoptotic cells, Q3. Annexin V-/PI+ indicate mechanically damaged cells, and Q4. Annexin V-/PI-indicate living cells. **e,** A representative micrograph of caspase 3/7 activity using CellEvent™ Green fluorescence indicate active caspase 3/7. **F,** Representative micrograph of apoptotic array proteins. **g,** Schematic for Z-VAD-FMK treatment to cells (10μM) and mice (5mg/kg). **h,** A graph showing percent PI stained population of Z-VAD-FMK treated and HSV-1 infected iRic KO MEFs. **i,** A Survival graph showing probability of survival for Z-VAD-FMK or mock treated HSV-1 infected iRic -/- animals.

To study the apoptotic responses of iRic -/- mice *in vitro*, we isolated mouse embryonic fibroblasts (MEFs) from murine foetuses and treated them with 4-hydroxytamoxifen (Fig. 3c). Thus, we were able to culture iRic +/+ and -/- cells for *in vitro* experiments. In accordance with our *in vivo* results, we found that the iRic -/- MEFs showed increased levels of both early and late stage apoptosis (Fig. 3d). Additionally, caspase 3/7 activation was enhanced in the -/- MEFs, another confirmation of their differential cell death response (Fig. 3e). To further understand the factors contributing to the onset of apoptosis in iRic-/-, we analyzed relative levels of proteins in an apoptosis protein array (Fig. 3f and Extended data 3a). Interestingly, anti-apoptotic proteins myeloid leukemia cell differentiation protein 1 (MCL-1) was preferentially up-regulated in the iRic +/+ cells whereas several pro-apoptotic proteins including tumor necrosis factor receptor 1 (TNFR1), hypoxia inducing factor (HIF-1α) and HSP-27 were up-regulated in infected iRic -/- cells. MCL-1 is known to attenuate apoptosis and promote survivalof neuronal cells^17^. Reduced levels of MCL-1 in the iRic -/- MEFs verifies the importance of Rictor in stabilizing the protein^18^. The cumulative evidence suggests that mTORC2 acts to inhibit the apoptotic response. Suppression of MCL-1 correlates with upregulation of PUMA which is under transcriptional regulation of Forkhead box O3a (FOXO3a)^19^.

If loss of Rictor results in an increase in cell deathupon viral infection both *in vitro* and *in vivo*, then an inhibitor ofapoptosis may promote host cell survival and prevent the lethality of infection in iRic -/- mice. We used the pan-caspase inhibitor Z-VAD-FMK to pharmacologically reduce apoptosis in both the iRic -/- MEFs and mice (Fig. 3g). Z-VAD-FMK treatment resulted in a loss of cell death in the MEFs (Fig. 3h). Critically, Z-VAD-FMK rescued the iRic -/- mice as 60% of the mice survived to 15 dpi as opposed to only 20% in the mock-treated group (Fig. 3i). Thus, loss of Rictor promotes exacerbated host apoptotic responses during infection that results in the death of the host *in vivo*.

After observing the extensive cell death responses in iRic -/- MEFs, we wanted to elucidate the mechanism of the observed apoptosis. mTORC2 has been shown to phosphorylate AKT at Ser-473 which activates it^20^. AKT then proceeds to phosphorylate the pro-apoptotic transcription factor Forkhead box O3a (FoxO3a)^21^ at Ser-253 and Thr-32. p-FoxO3a is inactive and precluded from the nucleus which prevents it from stimulating apoptosis (Fig. 4a)^21^. A western blot time course of infected HCE cells revealed that phosphorylation of AKT at 8 hpi is closely followed by phosphorylation of FoxO3a at 10 hpi (Fig. 4b and Extended data 4a). A similar phenomenon occurs in Lund human mesencephalic (LUHMES) neuronal cells whereby both AKT and FoxO3a become phosphorylated by 24 hpi, highlighting the mTORC2-Akt-FoxO3a axis as a conserved response during HSV-1 infection (Fig. 4c). We hypothesized that loss of Rictor would inhibit AKT phosphorylation which would allowFoxO3a to facilitate the transcription of pro-apoptotic factors. We found that deletion of Rictor abrogated phosphorylation of both AKT and FoxO3aduring viral infection (Fig. 4d). In contrast, the phosphorylation of both proteins occurred in a dose-dependent manner with infection in iRic +/+ MEFs (Fig. 4d). If the phosphorylation of FoxO3a is inhibited, it should be present inside of the nucleus and vice versa. Using confocal microscopy, we found that FoxO3a was precluded from the nucleus during HSV-1 infection in iRic +/+ MEFs (Fig. 4e). However, FoxO3a appeared inside of the nucleus in the -/- MEFs, localizing within the DAPI stain (Fig. 4e). Thus, loss of Rictor prevents the downstream phosphorylation of AKT and FoxO3a which stimulates cellular apoptosis.

**Figure 4.**
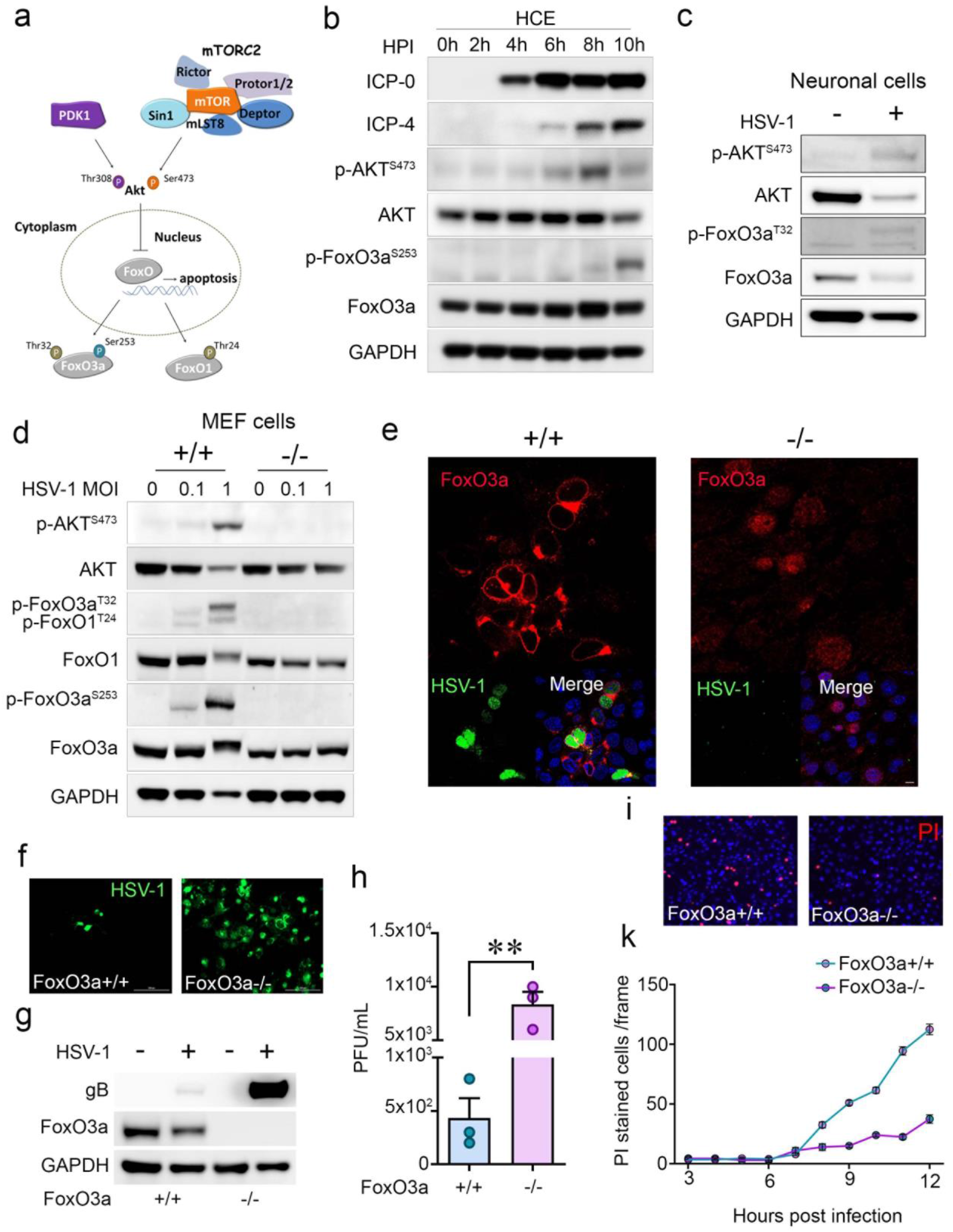
Rictor/mTORC2 dependent nuclear preclusion of FOXO3a is important for cell survival during viral infection. **a,** Schematic representation of the hypothesis. **b,** Representative western blots showing expression and phosphorylation of respective proteins at different hpi **c,** Representative western blot illustrating expression and phosphorylation of respective proteins in LUHMES cells. **d,** Representative western blot illustrating expression and phosphorylation of respective proteins in iRic MEF’s. **e,** Representative micrograph of immunofluorescence confocal imaging illustrating location of FoxO3a, Scale bar-10μm. **f,** Representative micrograph of fluorescence imaging in HSV-1 (GFP tagged) infected cells, Scale bar-200μm **g,** Representative micrograph of western blot showing indicated protein expression. **h,** Plaque assay showing PFU/mL. **I,** A representative micrograph showing PI stained population of HSV-1 infected MEFs at 12 hpi. **j,** A line graph showing PI stained population of HSV-1 infected MEFs.

We hypothesized that loss of FoxO3a reduces the cell’s ability to mitigate infection via apoptosis. By using cells with a genetic deletion of FoxO3a, we observed an increase in viral infection in the FoxO3a -/- cells as compared to the FoxO3a +/+ cells (Fig. 4f-h). Consistent with this result, FoxO3a -/- cells demonstrate slower apoptosis kinetics during infection (Fig. 4i,k). iRic -/- cells conversely show an increased rate of cell death during HSV-1 infection or treatment with the agonist etoposide, a pharmacological promoter of apoptosis (Extended Data Fig. 5a-d). The results are in congruent with our hypothesis that during HSV-1 infection nuclear preclusion of FOXO3a mediated through mTORC2-AKT axis is prerequisite for cell survival.

Collectively, our findings suggest that during HSV-1 infection, loss of Rictor prevents the mTORC2 complex from phosphorylating Akt. Inactive Akt cannot phosphorylate FoxO3a which allows it to stimulate apoptosis. Mice with infected brain stems experience an increase in neuronal cell death, which becomes lethal over time. In light of this information, we propose that mTORC2 is essential to control HSV-1 infection and cell survival during stress situation 1) by inducing innate and adaptive immune responses and 2) protecting host cells and neurons from stress-induced apoptosis. During herpes simplex encephalitis, neuronal apoptosis may contribute to severe symptoms such as hallucinations, partial paralysis, memory loss and even death. Acyclovir treatment is usually given well after the virus has reached the brain. However alongside of acyclovir, apoptosis inhibitor like Z-VAD-FMK may support mTORC2’s functions in host survival and serve as an adjuvant therapy for severe HSV-1 infections, reducing neurodegeneration during encephalitis. Furthermore, potentiating the pro-survival and antiviral activities of mTORC2 may comprise a novel treatment for severe cases of viral infections.

## Materials & Methods

### Cells and viruses

Human corneal epithelial (HCE) cell line (RCB1834 HCE-T) was procured from Kozaburo Hayashi (National Eye Institute, Bethesda, MD). HCE’s were cultured in minimum essential medium (MEM) (Life Technologies, Carlsbad, CA) supplemented with 10% fetal bovine serum (FBS) (Life Technologies) and 1% penicillin/streptomycin (Life Technologies).

Mouse embryonic fibroblasts (MEFs) with inducible RICTOR -/- were isolated from a mice having Rictor conditional alleles able to transiently express tamoxifen-inducible Cre recombinase (CreERT2). RICTOR knockout in the cells was achieved by treating them with 4-hydroxytamoxifen (4OHT) (2 μM) for 72h. MEFs with FoxO4^+/-^, p53^-/-^ immortalized FoxO3a+/+ and FoxO3a-/- were a generous gift from Prof. Nissim Hay (University of Illinois at Chicago, USA). DMEM containing 10% FBS and 1% penicillin/streptomycin was used to grow all MEFs. Different strains of HSV-1 used in study were KOS-WT, McKrae strains, both were provided by Dr. Patricia G. Spear (Northwestern University, Chicago, IL whereas Dual color ICP0p-GFP/gCp-RFP was gifted by Dr. Paul Kinchington (University of Pittsburgh). HSV-1 17 strain was obtained from Dr. Richard Thompson (University of Cincinnati, Cincinnati, OH). The virus stocks were made in Vero cells and stored at −80°C.

### Inducible Rictor knock out animal generation

Rictor^F/F^ mutant mice possessing loxP sites flanking exon 11 of the RPTOR independent companion of MTOR, complex 2 (Rictor) gene (STOCK Rictortm1.1Klg/SjmJ) and whole body cre/ERT2 mice (B6.129-Gt(ROSA)26Sortm1(cre/ERT2)Tyj/J) were purchased from The Jackson Laboratory. The mice were bred to create a inducible Rictor knockout mice (iRic-/-). In order to knock out Rictor from iRic-/- animals, they were treated with tamoxifen (2mg/kg) for five days and then used for the experiment.

### Antibodies

HSV-1 viral proteins were detected by ICP0 (ab6513), gB (ab6506) both purchased from Abcam (Cambridge, United Kingdom. Following antibodies were used for western blot or immunofluorescence were FoxO3a (2497S), AKT (9272S), p-FoxO3a^S253^ (9466S), RICTOR (2114S), p-FoxO1^T24^/p-FoxO3A^T32^ (9464S), p-AKT^S473^ (9271S), FoxO1 (FoxO1), Histone (4499S), IRF7 1:500 (4920) all purchased from cell signaling technology (Danvers, MA), ICP0 (sc-53070) and ICP4 (sc-69809) both purchased from Santa Cruz Biotechnology, (Dallas, TX) and GAPDH (10494-1-AP) was purchased from Proteintech Group, Inc., (Rosemont, IL).

### Chemical reagents

Pharmacological inhibitors including 4-hydroxytamoxifen (S7817), Z-VAD-FMK (S7023), Etoposide (S1225) were purchased from Selleckchem (Houston, TX).

### Western blot

The cells in a reaction well were collected using Hanks Cell Dissociation buffer. Protein lysates were prepared using radioimmunoprecipitation assay (RIPA) buffer (Sigma Aldrich, St Louis, MO) as per manufacturer’s guidelines. The protein samples were mixed with LDS sample loading buffer at 4X concentration followed by addition of beta-mercaptoethanol (5%) (Bio-Rad, Hercules, CA). The resultant mixture was denatured at 95°C for 9 min. The protein samples were then electrophoresed on Invitrogen™ Mini Gel Tank (Fisher scientific) through pre-casted gels (4-12%). The proteins were blotted on nitrocellulose membrane (Fisher scientific). The membrane was blocked in 5% milk/TBS-T for 1 h followed by overnight incubation with primary antibody at 4°C. The membranes were washed with TBS-T and incubated with respective horseradish peroxidase (HRP) conjugated secondary antibody (anti-mouse 1:10000 or anti-Rabbit 1:10000) for 1h at room temperature. 1:2500 concentrations of anti-rabbit were used for detection of all phosphoproteins. The membranes were washed again before exposing them to SuperSignal West Femto maximum sensitivity substrate (Thermo Scientific, Waltham, MA) and proteins were visualized using the with Image-Quant LAS 4000 biomolecular imager (GE Healthcare Life Sciences, Pittsburgh, PA).

### Quantitative real time polymerase chain reaction

RNA isolation was performed using TRIzol (Life Technologies). High Capacity cDNA Reverse Transcription kit (Life Technologies) was used to transcribe RNA to cDNA using High Capacity RNA–to-cDNA Kit (Applied Biosystems Foster City, CA). The cDNA was further used for Real-time quantitative polymerase chain reaction (qPCR) performed on QuantStudio 7 Flex system (Invitrogen™ Life Technologies) using Fast SYBR Green Master Mix (Life Technologies).

The primers used in this study are.

Glycoprotein D qPCR forward: 5’ - TACAACCTGACCATCGCTTC-3’

Glycoprotein D qPCR reverse: 5’ - GCCCCCAGAGACTTGTTGTA-3’

ICP-0 qPCR forward: 5’- ACAGACCCCCAACACCTACA-3’

ICP-0 qPCR reverse: 5’ - GGGCGTGTCTCTGTGTATGA-3’

Glycoprotein B qPCR forward: 5’ - GCCTTTTGTGTGTGTGTGGG-3’

Glycoprotein B qPCR reverse: 5’ - GCCTTTTGTGTGTGTGTGGG-3’

Human GAPDH qPCR forward: 5’ - TCCACTGGCGTCTTCACC-3’

Human GAPDH qPCR reverse: 5’ - GGCAGAGATGATGACCCTTTT-3’

Human RICTOR qPCR forward: 5’ - TGGGTGTGAACCATGAGAAGTATG-3’

Human RICTOR qPCR reverse: 5’ - GGTGCAGGAGGCATTGCT-3’

Mouse interferon-α qPCR forward: 5’-CCTGCTGGCTGTGAGGAAAT-3’

Mouse interferon-α qPCR reverse: 5’-GACAGGGCTCTCCAGACTTC-3’

Mouse interferon-β qPCR forward: 5’- TGTCCTCAACTGCTCTCCAC-3’

Mouse interferon-β qPCR reverse: 5’- CATCCAGGCGTAGCTGTTGT-3’

Mouse interleukin-6 qPCR forward: 5’-ACGGCCTTCCCTACTTCACA-3’

Mouse interleukin-6 qPCR reverse: 5’-CATTTCCACGATTTCCCAGA-3’

Mouse interleukin-12 qPCR forward: 5’-AAATGAAGCTCTGCATCCTGC-3’

Mouse interleukin-12 qPCR reverse: 5’-TCACCCTGTTGATGGTCACG-3’

Mouse TNF-α qPCR forward: 5’-GCCTCTTCTCATTCCTGCTTG-3’

Mouse TNF-α qPCR reverse: 5’-CTGATGAGAGGGAGGCCATT-3’

Mouse β-actin qPCR forward: 5’- CGGTTCCGATGCCCTGAGGCTCTT-3’

Mouse β-actin qPCR reverse: 5’- CGTCACACTTCATGATGGAATTGA-3’

### Plaque assay

Either HCE or MEFs were infected with either mock or 0.1 MOI of HSV-1 and incubated at 37°C, 5% CO2. After 2hpi the inoculum medium was replaced by whole medium (MEM or DMEM containing 10% FBS and 1% PenStrap). The cells were incubated at 37°C, 5% CO_2_ for 24 h unless specified. After infection the cells were collected using Hanks buffer and suspended in Opti-MEM (Thermo Fisher Scientific). The resultant mixture was sonicated to produce lysates, which were used to quantify infectious virus particles. The VERO cells were seeded in tissue culture plate to form a monolayer. The monolayer was infected with respective dilution of cell lysate in Opti-MEM. At 2hpi the dilutions of cell lysate were aspirated and replaced by a complete medium (DMEM with 10% FBS and 1% PenStrap) containing 0.5% methylcellulose (Fisher Scientific). The cells were incubated at 37°C, 5% CO2, for 72 h. After incubation the cells were fixed by adding 100% methanol. After 20 mins of fixation the medium was removed and cells were stained with crystal violet solution. Numbers of plaques were counted visually.

### Immunofluorescence microscopy

HCE or MEFs were cultured in glass bottom dishes (MatTek Corporation, Ashland, MA). 24 hpi the cells were fixed with 4 % paraformaldehyde (PFA) and permeabilized with 0.1 % triton X-100 for 10 mins for intracellular staining with a mouse or rabbit monoclonal antibody against the target protein (1h) followed by a secondary antibody conjugated with FITC- or Alexa Fluor 647 (Sigma-Aldrich) (1.100) for 1h at room temperature. NucBlueLiveReady Probes Hoescht stain (Thermo R37605) was used to stain the nucleus (histone). The samples were investigated under 63X oil immersion objective using LSM 710 confocal microscope (Zeiss). The resulting images were processed using ZEN black software.

### Live cell imaging

iRic +/+ and iRic -/- MEFs were cultured in 12 well plate and infected with HSV-1 dual color fluorescent virus expressing GFP and RFP driven by ICP0 and gC promoter respectively. At 2 hpi, inoculation media was replaced with complete DMEM, and cells were placed in the incubation chamber of Zeiss spinning disk live-cell imaging system, which maintains 37°C and 5% CO_2_. Images were captured for RFP, GFP, and brightfield at an interval of 30 min for 24 h, and analyzed with ZEN software. Similar experiments were done for estimation of propidium iodide (PI) uptake for iRic +/+ and iRic -/- MEFs either infected with HSV-1 infection or treated with etoposide (10 μM). PI uptake experiment was done for HSV-1 infected FoxO3a +/+ and FoxO3a -/- MEFs as well. In all experiments NucBlueLiveReady Probes Hoescht stain was used to stain the nucleus (Thermo R37605).

### Infection to murine cornea

All animal care and procedures were performed in accordance with the institutional and NIH guidelines, and approved by the Animal Care Committee at University of Illinois at Chicago (ACC protocol 17-077). A C57BL/6J mice were either dosed with mock or tamoxifen (2mg/kg) for 5 days before infection. At the time of infection mice were anesthetized and their corneas were scarified in a 3×3 grid using a 30-gauge needle. The corneas were infected with HSV-1 McKrae (1 × 10^5^). At day 4 post infection eye wash samples were collected to quantify HSV-1 replication through plaque assay. Mice with acute weight loss (>20%) were euthanized for humane reasons. In a different set of experiment mice were checked for presence of HSV-1 replication in tissues like blood, trigeminal ganglion and brain stem. The apoptosis in ocular tissue and brain stem was analyzed by TUNEL staining as per manufacturer’s protocol (abcam inc.).

### Proteome profiler apoptosis array

iRic +/+ and iRic -/- cells were infected with HSV-1 at 1 MOI. After 12 hpi the cells were rinsed with PBS and the cells were lysed as per the manufacturer’s guidelines. The resultant mixture was centrifuged at 14,000 g and the supernatant was used to estimate the total protein by using Pierce™ BCA Protein Assay Kit (Thermo scientific). Equal quantity of protein (300 μg) was loaded onto each membrane. Detection of apoptosis related protein was further performed by the manufacturer’s guidelines (Proteome profiler™ array).

### Apoptosis detection by flow cytometry

iRic +/+ and iRic -/- cells were infected with HSV-1 at 1 MOI and harvested at 8 and 12 hpi. FITC Annexin V/Dead Cell Apoptosis Kit (Invitrogen™) was used to detect apoptosis. As per the manufacturer’s protocol the sample pellets were washed with cold PBS and suspended in 1x annexin binding buffer. The suspension was mixed with 5 μl FITC Annexin V and 1 μl of a propidium iodide solution (100 μg/mL) for 15 min in the cold. The mixture was diluted with 1X annexin binding buffer. Further flow cytometry (BD Accuri™ C6 Plus flow cytometer) was used to analyze the staining for Annexin V and propidium iodide. Flow data was analyzed using FlowJo software (Tree Star Inc.).

### Caspase3/7 activity assay

CellEvent™ Caspase3/7 green detection reagent was used for analyzing the activity of caspase 3/7 in iRic +/+ and iRic -/- cells were infected with HSV-1 at 1 MOI. After 12 hpi the cells were imaged using fluorescence microscopy. Caspase 3/7 activity in cells was identified by appearance of fluorescing bright green cells.

### Statistics Analysis

Error bars of all Figures represent SEM of three independent experiments (*n*=3), unless otherwise specified. The experimental dataset between the two groups have been compared using the two tailed unpaired Student’s t-test or 2way ANOVA. The *p* values have been added in Figures. Differences between values were considered significant when *P*=0.05.

### Key resource table

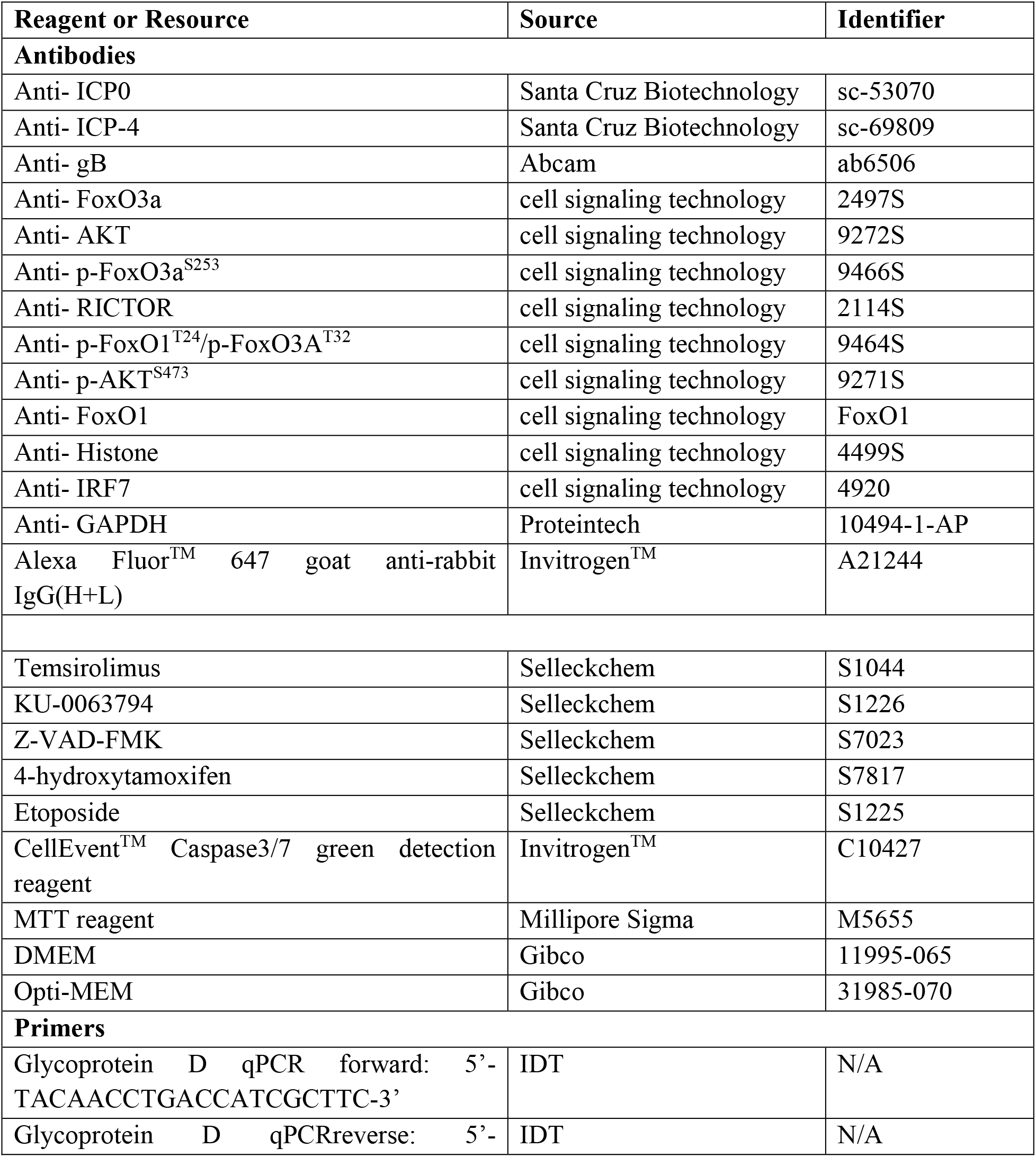
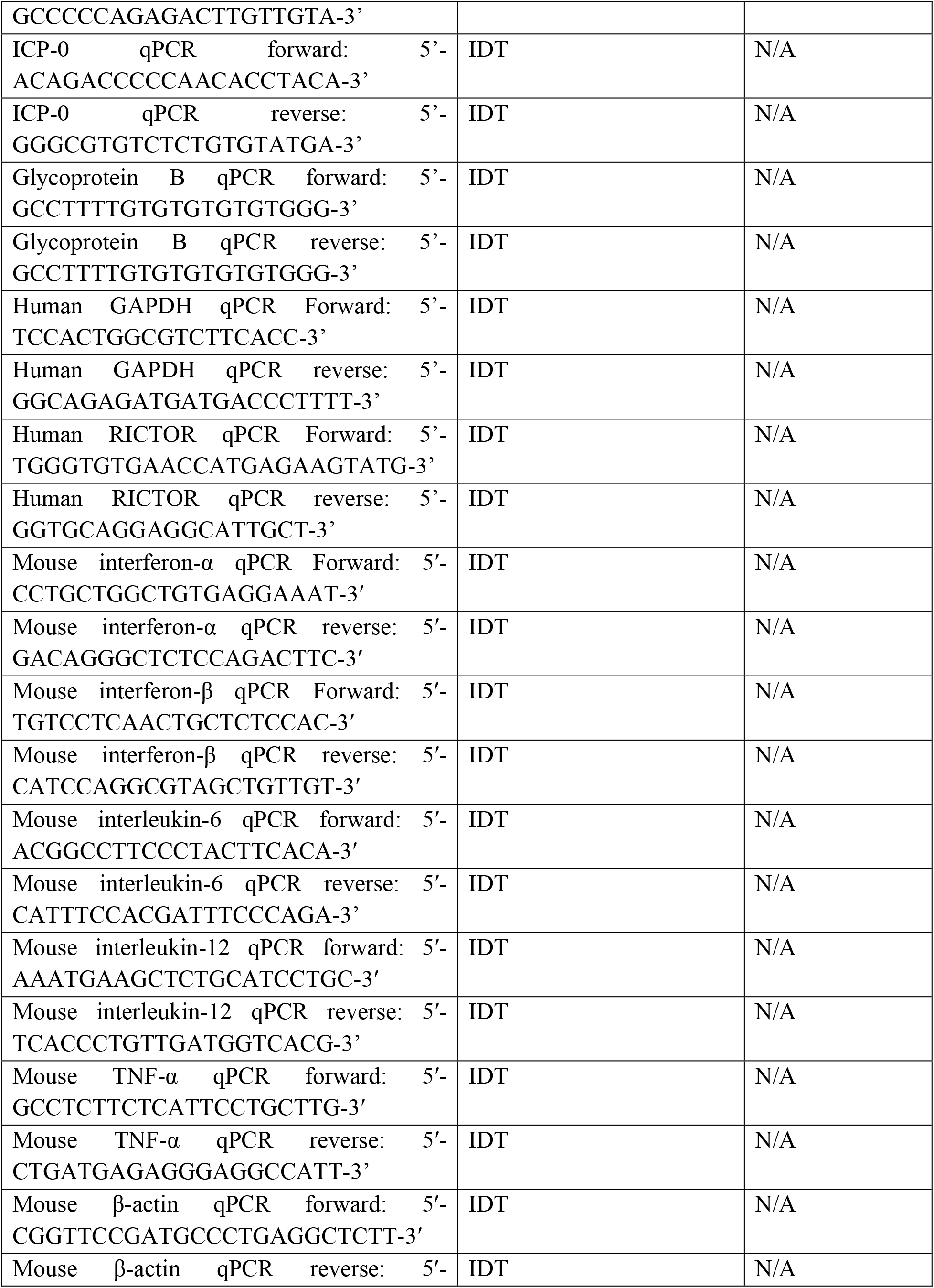
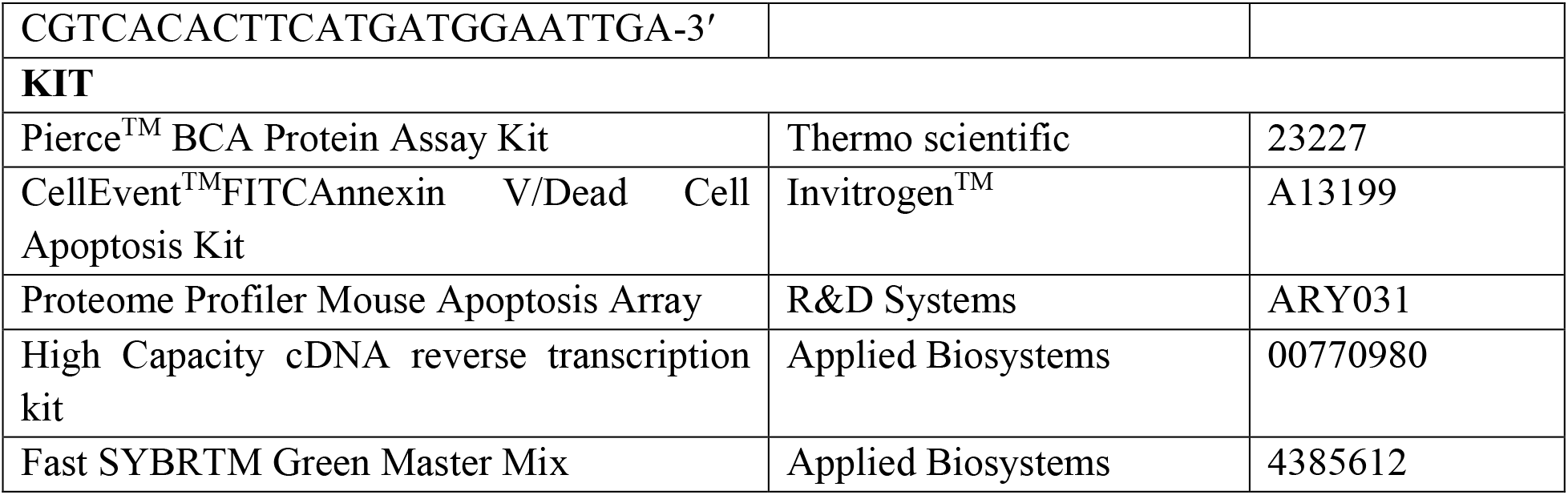

## Acknowledgements

We are grateful to Dr. Nissim Hay for FoxO3a^-/-^ cells. This work was supported by the National Institutes of Health RO1 grants EY029426, AI139768, EY024710 (to DS) and a NEI core grant (EY001792) and Illinois society for prevention of blindness.

## Author contributions

Conceptualization, R.K.S., and D.S.; Methodology, R.K.S., C.D.P., A.A, T.Y., J.M.A., L.K., K.R.; Investigation, R.K.S., C.D.P., A.A, D.S.; Writing Original Draft, R.K.S., and D.S.; Writing – Review and Editing, R.K., R.K., and D.S.; Funding Acquisition, R.K.S., D.S.; Supervision, D.S.

## References

1. Barber, G. N. Host defense, viruses and apoptosis. Cell Death Differ. 8, 113–126 (2001).

2. You, Y. et al. The suppression of apoptosis by α-herpesvirus. Cell Death Dis 8, e2749 (2017).

3. Deleidi, M. & Isacson, O. Viral and Inflammatory Triggers of Neurodegenerative Diseases. Sci Transl Med 4, 121ps3 (2012).

4. Kuss-Duerkop, S. K. et al. Influenza virus differentially activates mTORC1 and mTORC2 signaling to maximize late stage replication. PLoS Pathog. 13, e1006635 (2017).

5. Sato, R. et al. Combating herpesvirus encephalitis by potentiating a TLR3-mTORC2 axis. Nat. Immunol. 19, 1071–1082 (2018).

6. Shiota, C., et al. Multiallelic disruption of the rictor gene in mice reveals that mTOR complex 2 is essential for fetal growth and viability. Dev. Cell 11, 583–589 (2006).

7. Jullien, N. et al. Use of ERT2-iCre-ERT2 for conditional transgenesis. Genesis 46, 193–199 (2008).

8. Koganti, R., et al. Current and Emerging Therapies for Ocular Herpes Simplex Virus Type-1 Infections. Microorganisms 7 (2019).

9. Nicoll, M. P. et al. The HSV-1 Latency-Associated Transcript Functions to Repress Latent Phase Lytic Gene Expression and Suppress Virus Reactivation from Latently Infected Neurons. PLoS Pathog 12 (2016).

10. Lazorchak, A. & Su, B. Perspectives on the role of mTORC2 in B lymphocyte development, immunity and tumorigenesis. Protein Cell 2, 523–530 (2011).

11. Hao, Y. et al. The Kinase Complex mTOR Complex 2 Promotes the Follicular Migration and Functional Maturation of Differentiated Follicular Helper CD4+ T Cells During Viral Infection. Frontiers in immunology 9, 1127 (2018).

12. Wei, W. et al. Mammalian target of rapamycin complex 2 (mTORC2) regulates LPS-induced expression of IL-12 and IL-23 in human dendritic cells. J. Leukoc. Biol. 97, 1071–1080 (2015).

13. Raïch-Regué, D. et al. mTORC2 Deficiency in Myeloid Dendritic Cells Enhances Their Allogeneic Th1 and Th17 Stimulatory Ability after TLR4 Ligation In Vitro and In Vivo. J. Immunol. 194, 4767–4776 (2015).

14. Tenkerian, C. et al. mTORC2 Balances AKT Activation and eIF2α Serine 51 Phosphorylation to Promote Survival under Stress. Mol Cancer Res 13, 1377–1388 (2015).

15. Wen, Y., et al. mTORC2 activation protects retinal ganglion cells via Akt signaling after autophagy induction in traumatic optic nerve injury. Experimental & molecular medicine 51, 1–11 (2019).

16. Luo, H. R. et al. Akt as a mediator of cell death. Proc Natl Acad Sci U S A 100, 11712–11717 (2003).

17. Arbour, N. et al. Mcl-1 Is a Key Regulator of Apoptosis during CNS Development and after DNA Damage. J Neurosci 28, 6068–6078 (2008).

18. Koo, J., et al. mTOR Complex 2 Stabilizes Mcl-1 Protein by Suppressing Its Glycogen Synthase Kinase 3-Dependent and SCF-FBXW7-Mediated Degradation. Molecular and cellular biology 35, 2344–2355 (2015).

19. Li, H., et al. Downregulation of MCL-1 and upregulation of PUMA using mTOR inhibitors enhance antitumor efficacy of BH3 mimetics in triple-negative breast cancer. Cell Death Dis 9, 137 (2018).

20. Manning, B. D. & Toker, A. AKT/PKB Signaling: Navigating the Network. Cell 169, 381–405 (2017).

21. Zhang, X., et al. Akt, FoxO and regulation of apoptosis. Biochim. Biophys. Acta 1813, 1978–1986 (2011).

