## Supplement file for "mTORC2 combats cellular stress and potentiates immunity during viral infection"

### Summary
Herpes simplex virus type 1 (HSV-1) causes ocular and orofacial infections, which are generally well controlled by the host and nonlethal. In rare cases, HSV-1 causes encephalitis, which leads to permanent brain injuries, memory loss or even death. Host factors protect the organism from viral infections by activating the immune response. However, the factors that confer neuroprotection during viral encephalitis are unknown. Here we show that mammalian target of rapamycin complex 2 (mTORC2) is essential for the host survival of ocular HSV-1 infections *in vivo*. We found that the loss of mTORC2 causes systemic HSV-1 infection not only because of weak innate and adaptive immune responses but also due to increased ocular and neuronal cell death, which becomes lethal over time. Furthermore, we found that mTORC2 mediates cell survival channels through the inactivation of the proapoptotic factor FoxO3a. Our results demonstrate how mTORC2 potentiates host defenses against viral infections as well as implicating mTORC2 as a necessary host factor for survival. We anticipate our findings may help develop new therapeutic window for severe HSV-1 infections, such as herpes simplex encephalitis.

**Keywords:** mTORC2, HSV-1, Systemic infection, immune response, apoptosis

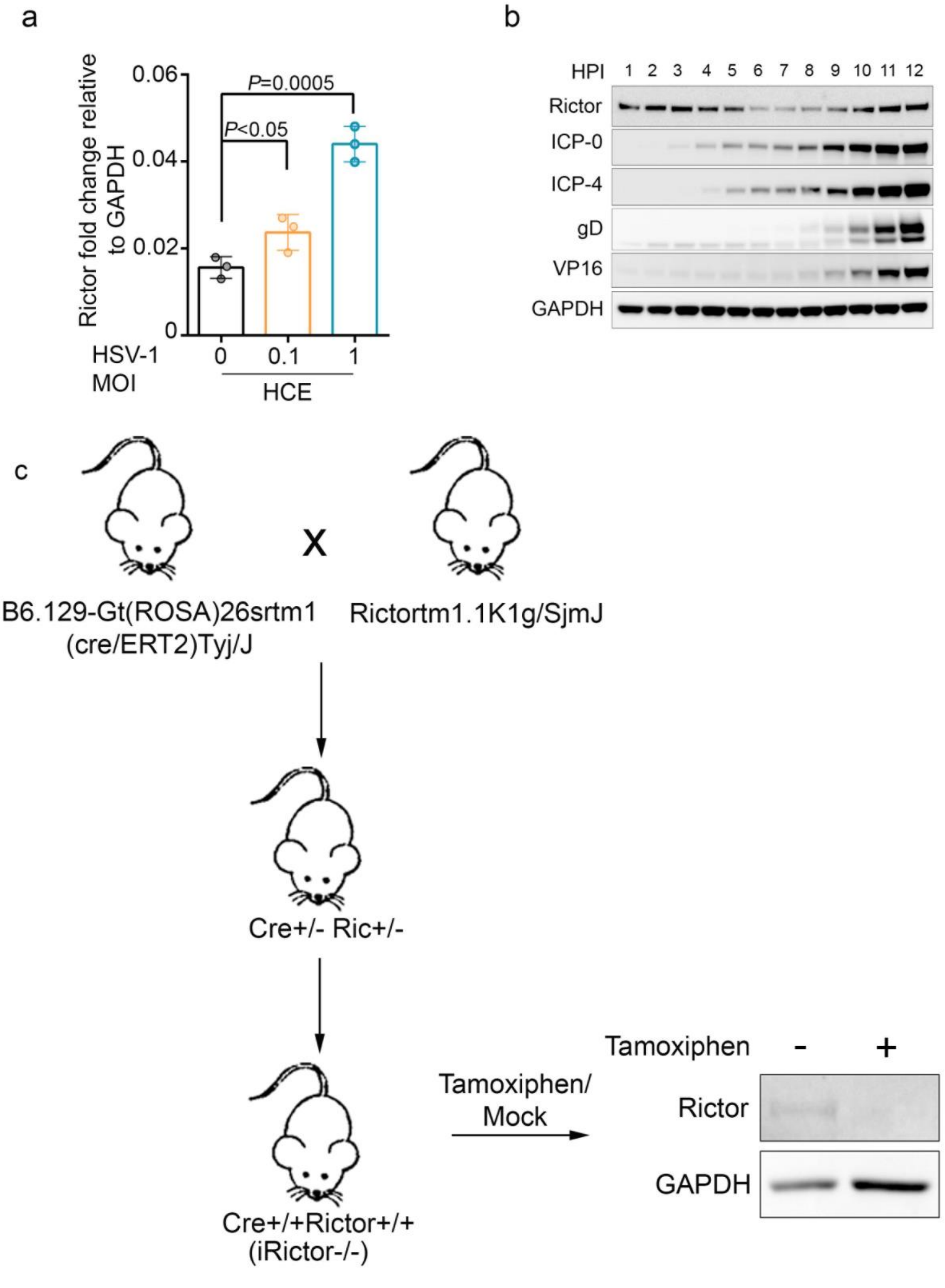


**Extended data 1| a.** qRTPCR analysis showing increase expression of Rictor with respect to HSV-1 infection in Human corneal epithelial cells. **b.** A representative image of protein expression in HSV-1 (1 multiplicity of infection) infected HCE cells at different time points. **c.** Schematic for the development of conditional rictor knockout mouse model.

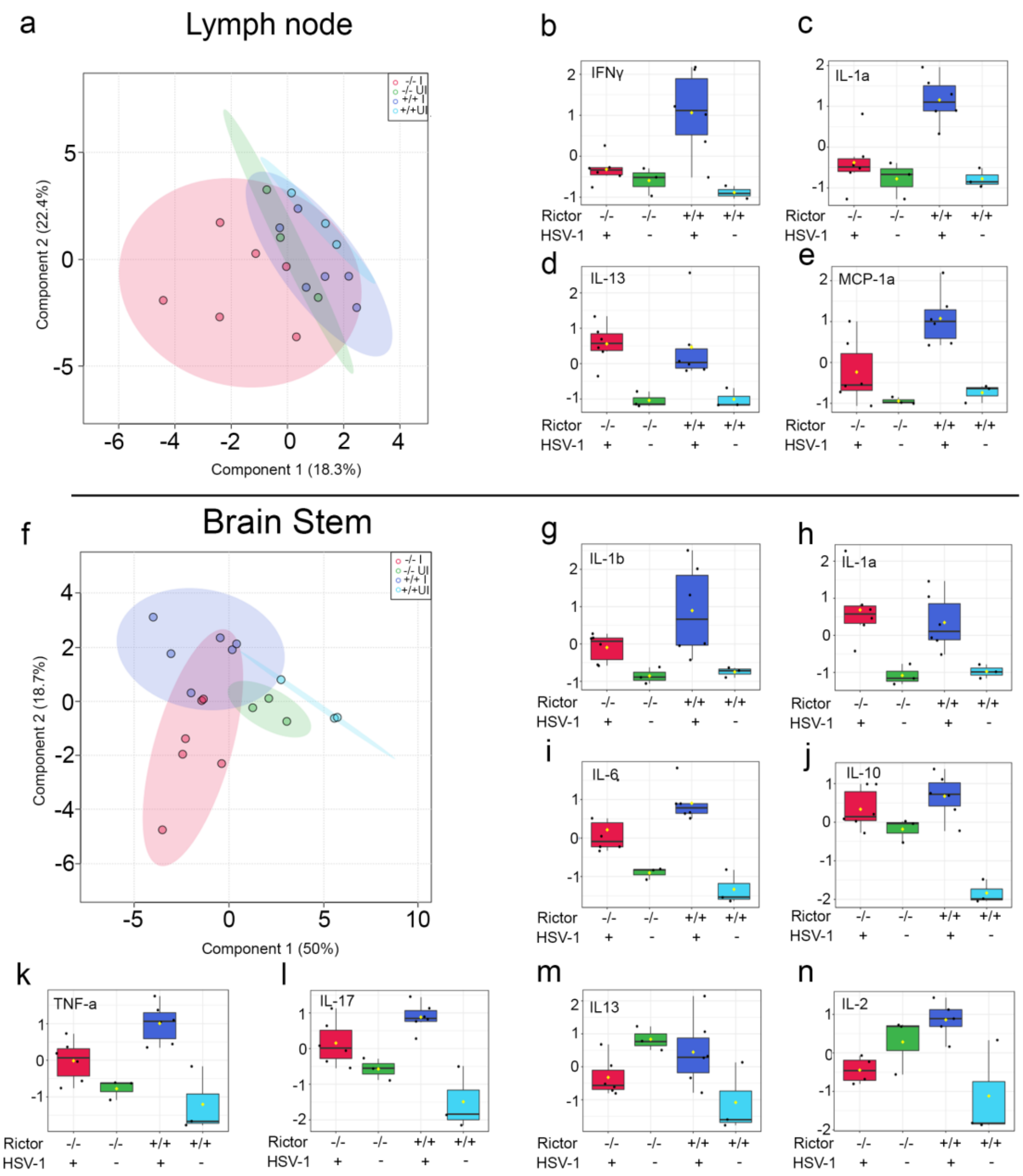


**Extended data 2| Cytokine levels in lymph node and brain stem tissue show differences in iRic +/+ or -/- mice a.** Principle component analysis of cytokine expression in lymph node tissue of HSV-1 infected mice **b-e.** A graph showing expression of respective cytokines in lymph node of HSV-1 or mock infected iRic +/+ or -/- mice. **f.** Principle component analysis of cytokine expression in brain stem tissue of HSV-1 infected mice **g-n.** A graph showing expression of respective cytokines in brain stem of HSV-1 or mock infected iRic +/+ or -/- mice.

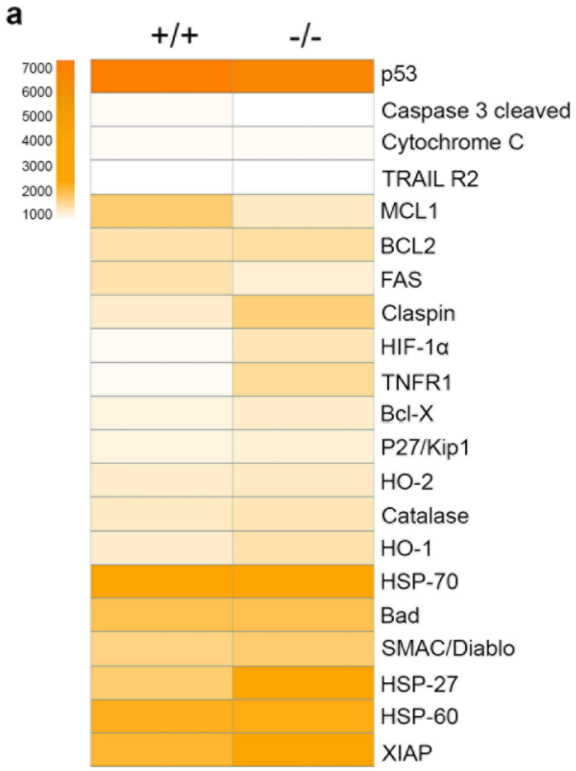


**Extended data 3| Differential expression of apoptosis related proteins in HSV-1 infected iRic +/+ or -/- cells a.** A heat map showing the quantitative analysis of proteome profiler mouse apoptosis array proteins from figure 3f.

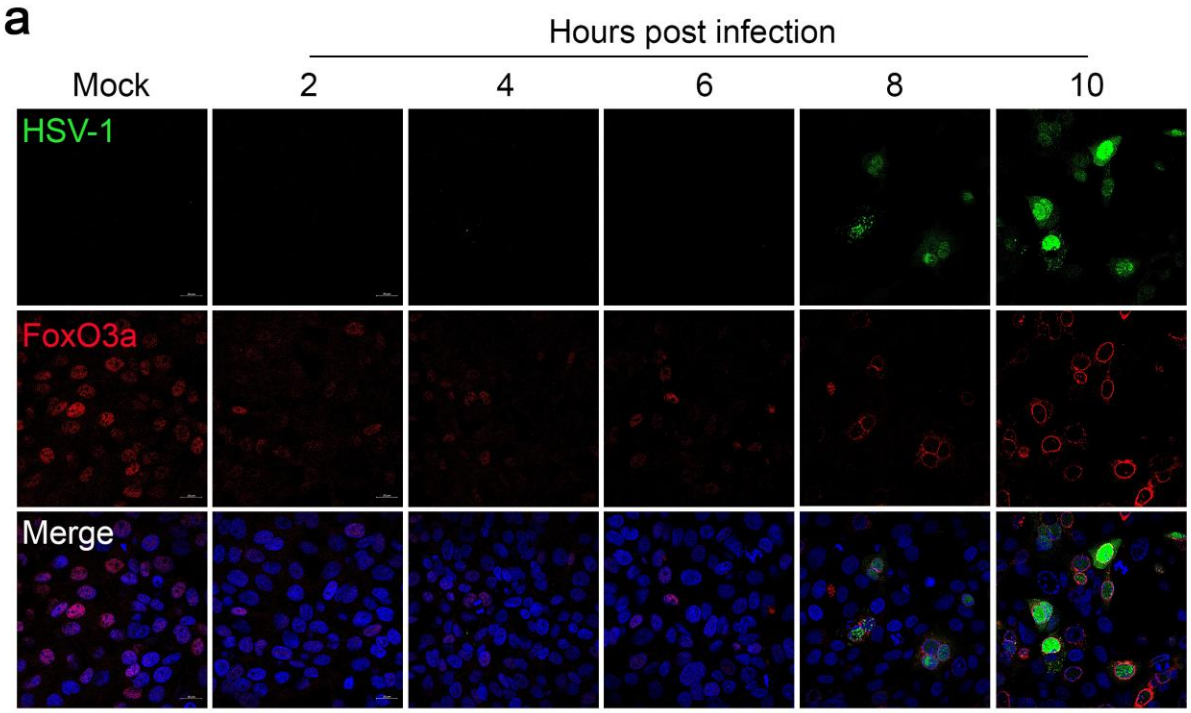


**Extended data 4| Time point analysis of FoxO3a localization in HSV-1 infected HCE a.** Representative micrograph of immunofluorescence confocal imaging illustrating expression and location of FoxO3a protein in HSV-1 infected HCE cells. Images were taken at different hpi.

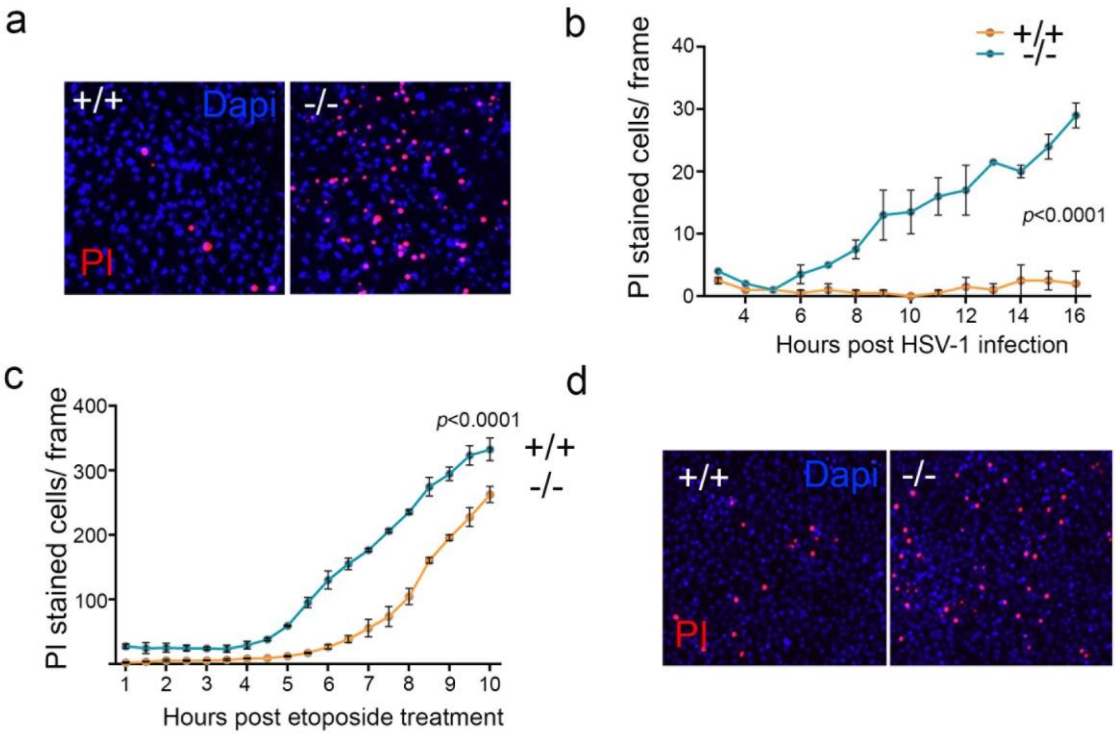


**Extended data 5| iRic-/- cells are sensitive to cell death with HSV-1 infection or etoposide treatment a.** Representative micrograph of fluorescent imaging for HSV-1 infected iRic +/+ or -/- MEFs stained with propidium iodide (red) and DAPI (blue) at 12hpi. **b.** Graph showing number of cells stained with PI at indicated time points. **c.** Graph showing number of iRic +/+ or -/- MEF cells treated with etoposide and stained with PI at indicated time points. **d.** Representative micrograph of time point fluorescent imaging of iRic +/+ or -/- MEFs treated with etoposide at 8h post treatment.
